# Therapeutic neutralizing monoclonal antibody administration protects against lethal Yellow Fever infection

**DOI:** 10.1101/2022.05.16.491863

**Authors:** Michael J. Ricciardi, Lauren N. Rust, Nuria Pedreño-Lopez, Sofiya Yusova, Sreya Biswas, Gabriela M. Webb, Lucas Gonsales-Nieto, Thomas B. Voigt, Johan J. Louw, Fernanda D. Laurino, John R. DiBello, Hans-Peter Raué, Aaron M. Barber-Axthelm, Samantha Uttke, Lidiane M.S. Raphael, Aaron Yrizarry-Medina, Brandon C. Rosen, Rebecca Agnor, Lina Gao, Caralyn Labriola, Michael Axthelm, Jeremy Smedley, Justin G. Julander, Myrna C. Bonaldo, Laura M. Walker, Ilhem Messaoudi, Mark K. Slifka, Dennis R. Burton, Esper G. Kallas, Jonah B. Sacha, David I. Watkins, Benjamin J. Burwitz

## Abstract

Few countermeasures to treat Yellow Fever virus (YFV) infection are under development, because vaccines have helped to limit new infections. Unfortunately, vaccine hesitancy, supply deficits, and a paucity of therapeutic options have left individuals at risk. Here, we tested potent YFV-specific neutralizing monoclonal antibodies in rodents and non-human primates. We administered antibodies during acute pathogenic YFV infection and demonstrate that we can prevent severe disease and death. Given the severity of YFV-induced disease, our results show that these antibodies could be effective in saving lives and fill a much-needed void in managing Yellow Fever cases during outbreaks around the world.

**One Sentence Summary:** Therapeutic monoclonal antibodies prevent death from YFV infection.

## INTRODUCTION

Yellow Fever virus (YFV) infection results in high viral loads and a mortality rate of up to 50% of hospitalized patients (*1*). Even with the availability of the live-attenuated YFV-17D vaccine, the World Health Organization (WHO) estimates an annual incidence of 200,000 YFV cases, with 29,000 to 60,000 related deaths (*2*). Despite numerous vaccine campaigns, immunization coverage remains low, resulting in a significant number of at-risk individuals with the potential for international spread (*3*). This is exacerbated by the uncommon (1 in 250,000) but severe side-effects caused by the YFV-17D vaccine, which can lead to severe illness, organ failure, or death (*4*). During the 2016-2019 outbreak in Brazil, the WHO reported approximately 100 cases of severe adverse events due to mass vaccination campaigns, which dissuaded many people from receiving the vaccine (*3, 5*).

There is currently no approved treatment for YFV infection, and there are no active clinical trials registered on clinicaltrials.gov testing interventional YFV therapeutics. Previous clinical trials have included two limited studies. A Phase Ib clinical trial (NCT03891420), which compared intravenous infusion of the adenosine nucleoside analogue Galidesivir against placebo in patients hospitalized for YFV infection, was terminated by the sponsor early in April 2021 due to a lack of cases. Another Phase Ia/b clinical trial (NCT03776786) studied the safety, pharmacokinetics, and efficacy of an intravenously administered human monoclonal antibody (mAb) TY104. The sponsors found that the mAb infusions were well tolerated and efficacious; however, the study faced data limitations, only measuring abrogation of viremia following vaccination with YFV-17D, an attenuated YFV vaccine strain (*6*). Thus, no clinical data exists showing successful drug interventions for severe YFV infection.

Our collaborators at Adimab recently published over 1,200 YFV-specific mAb sequences from YFV-17D vaccinated volunteers (*7*). Here, we further characterize a subset of these human IgG1 mAbs for their ability to neutralize lab-adapted and primary isolates of YFV both *in vitro* and *in vivo*. We show that several mAbs successfully neutralize primary isolates of YFV obtained from patients in Brazil at half-maximal inhibitory concentration values (IC_50_) of less than 50 ng/mL. In addition, we show that passive infusion of two high-avidity, YFV-specific mAbs can ameliorate YFV disease in both Syrian golden hamsters and Indian rhesus macaques (RM), providing a strong rationale for expedited clinical development of this intervention.

## RESULTS

### Antibody screening

We began by screening 489 YFV-specific mAbs isolated from memory B-cells of YFV-17D-immunized volunteers for neutralization against YFV-17D (*7*). From this analysis, we identified 38 mAbs that neutralized YFV-17D with an IC**_50_** < 200 ng/mL (Table 1). Only one of these mAbs also showed binding to other tested flaviviruses which included Zika, Dengue, and West Nile virus. Four of these mAbs targeted YFV envelope domain III (DIII), thirty targeted YFV envelope domain II (DII), and four others were unable to be clearly mapped. All DII mAbs outcompeted another previously isolated DII mAb 5A, which showed protection in a mouse YFV challenge study (*8*). The mAbs were then further down-selected by assessing their ability to neutralize other strains of YFV, specifically a primary wild-type isolate (YFV-ES504) from the 2016-2017 outbreak in Brazil (*9*). Of the 38 mAbs, only 16 neutralized YFV-ES504 at an IC_50_ < 1 μg/mL. Of note, only five mAbs neutralized 100% of YFV-ES504 at < 500 ng/mL. Synergy studies were also performed with these five mAbs, however, no additive synergy was observed (data not shown).

**Table 1.** Summary of screened mAbs.

| mAb ID | YFV-E Domain | YFV-17D IC<sub>50</sub> (ng/ml) | YFV-ES504 IC<sub>50</sub> (ng/ml) | YFV-ES504 Vmax (ng/ml) | VH usage | VL usage |
| --- | --- | --- | --- | --- | --- | --- |
| MBL-YFV-01 | DII | 2.6 | <31 | 250 | IGHV3-30 | IGLV1-51 |
| MBL-YFV-02 | DIII | 0.9 | <31 | <31 | IGHV3-33 | IGLV1-44 |
| MBL-YFV-03 | DII | 8.5 | >1,000 | >1,000 | IGHV4-4 | IGLV1-51 |
| MBL-YFV-04 | DII | 25.6 | >1,000 | >1,000 | IGHV3-11 | IGLV2-14 |
| MBL-YFV-05 | DII | 90.1 | >1,000 | >1,000 | IGHV4-61 | IGKV3-11 |
| MBL-YFV-06 | DII | 15.2 | >1,000 | >1,000 | IGHV4-4 | IGLV1-51 |
| MBL-YFV-07 | DII | 22.2 | >1,000 | >1,000 | IGHV3-33 | IGKV1D-39 |
| MBL-YFV-08 | DII | 6.5 | >1,000 | >1,000 | IGHV4-4 | IGLV1-51 |
| MBL-YFV-09 | DII | 2.5 | >1,000 | >1,000 | IGHV4-38-2 | IGLV1-51 |
| MBL-YFV-10 | DII | 50.1 | <31 | >1,000 | IGHV4-4 | IGLV1-51 |
| MBL-YFV-11 | DII | 16.2 | >1,000 | >1,000 | IGHV4-4 | IGLV1-51 |
| MBL-YFV-12 | DII | 42.9 | 930 | >1,000 | IGHV4-4 | IGLV1-51 |
| MBL-YFV-13 | DII | 14.2 | >1,000 | >1,000 | IGHV4-4 | IGLV1-51 |
| MBL-YFV-14 | DII | 182.1 | >1,000 | >1,000 | IGHV4-61 | IGKV3-11 |
| MBL-YFV-15 | DII | 147.9 | >1,000 | >1,000 | IGHV3-11 | IGLV3-21 |
| MBL-YFV-16 | DII | 107.5 | >1,000 | >1,000 | IGHV3-11 | IGLV1-51 |
| MBL-YFV-17 | DIII | 38.1 | 340 | 1,000 | IGHV3-23 | IGKV1-33 |
| MBL-YFV-18 | DII | 7.4 | 50 | 1,000 | IGHV4-4 | IGLV1-51 |
| MBL-YFV-19 | DIII | 3.0 | <31 | 125 | IGHV3-33 | IGLV1-44 |
| MBL-YFV-20 | DIII | 10.4 | <31 | 1,000 | IGHV3-23 | IGKV3D-15 |
| MBL-YFV-21 | DII | 175.2 | >1,000 | >1,000 | IGHV3-11 | IGLV3-21 |
| MBL-YFV-22 | Unk | 12.1 | >1,000 | >1,000 | IGHV3-30 | IGLV3-21 |
| MBL-YFV-23 | Unk | 3.1 | <31 | 1,000 | IGHV3-9 | IGLV6-57 |
| MBL-YFV-24 | Unk | 92.7 | 1,000 | >1,000 | IGHV3-74 | IGLV1-51 |
| MBL-YFV-25 | DII | 44.9 | >1,000 | >1,000 | IGHV3-11 | IGLV1-51 |
| MBL-YFV-26 | DII | 23.6 | >1,000 | >1,000 | IGHV4-4 | IGLV1-51 |
| MBL-YFV-27 | DII | 13.9 | >1,000 | >1,000 | IGHV4-4 | IGLV1-51 |
| MBL-YFV-28 | DII | 4.3 | 40 | >1,000 | IGHV3-23 | IGKV1-39/P |
| MBL-YFV-29 | DII | 2.2 | 1,000 | 1,000 | IGHV4-38-2 | IGLV1-51 |
| MBL-YFV-30 | DII | 6.4 | >1,000 | >1,000 | IGHV4-4 | IGLV1-51 |
| MBL-YFV-31 | DII | 3.2 | 1,000 | >1,000 | IGHV3-11 | IGLV2-14 |
| MBL-YFV-32 | DII | 20.8 | 30 | 500 | IGHV3-30 | IGKV4-1 |
| MBL-YFV-33 | DII | 8.8 | 1,000 | >1,000 | IGHV4-4 | IGLV1-51 |
| MBL-YFV-34 | DII | 5.3 | >1,000 | >1,000 | IGHV3-11 | IGLV2-14 |
| MBL-YFV-35 | Unk | 8.8 | >1,000 | >1,000 | IGHV3-30 | IGKV1-33 |
| MBL-YFV-36 | DII | 24.1 | <31 | <31 | IGHV3-11 | IGKV3D-15 |
| MBL-YFV-37 | DII | 64.8 | >1,000 | >1,000 | IGHV3-30 | IGLV2-14 |
| MBL-YFV-38 | DII | 13.2 | >1,000 | >1,000 | IGHV4-59 | IGKV3-11 |

Unk = Unknown epitope
IC<sub>50</sub> = Concentration of mAb to neutralize 50% of the viral plaques
Vmax = Minimum mAb concentration to neutralize 100% of the viral plaques

We selected two final antibodies (MBL-YFV-01 and MBL-YFV-02) that did not compete with each other and neutralized 100% of YFV-ES504 at < 125 ng/mL, then assessed their capacity to neutralize three additional primary isolates from Brazil (YFV-4408-1E, YFV-RJ155, and YFV-GO09) (Table 2). These strains were chosen because they were isolated during different outbreaks (*10*). YFV-4408-1E was isolated from a YFV-infected howler monkey in 2008 in Rio Grande do Sul state. YFV-RJ155, similar to the YFV-ES504 strain, was obtained from the most recent outbreak in southeastern Brazil (2016-2017). YFV-GO09 was isolated in Goiás (2017), yet it was not related to the 2016-2017 outbreak. Both mAbs had an IC_50_ < 15 ng/mL against all of these primary, highly pathogenic isolates.

**Table 2.** Broadly neutralizing mAbs.

| mAb ID | YFV-17D IC<sub>50</sub><br>(ng/mL) | YFV-Jimenez<br>IC<sub>90</sub> (ng/mL) | YFV-DakH1279<br>IC<sub>50</sub> (ng/mL) | YFV-ES504<br>IC<sub>50</sub> (ng/mL) | YFV-ES504<br>Vmax (ng/mL) | YFV-4408-1E<br>IC<sub>50</sub> (ng/mL) | YFV-RJ155<br>IC<sub>50</sub> (ng/mL) | YFV-GO09<br>IC<sub>50</sub> (ng/mL) |
| --- | --- | --- | --- | --- | --- | --- | --- | --- |
| MBL-YFV-01 | 0.9 | 2 | 75 | <15 | <31 | <15 | <15 | <15 |
| MBL-YFV-02 | 2.6 | 11 | 75 | <15 | 125 | <15 | <15 | <15 |

IC50 = Concentration of mAb to neutralize 50% of the viral plaques
IC90 = Concentration of mAb to neutralize 90% of the viral plaques
Vmax = Minimum mAb concentration to neutralize 100% of the viral plaques

In further preparation for *in vivo* experiments, we evaluated both mAbs against YFV-Jimenez, a Mexican isolate used in the Syrian golden hamster models (*11*), and YFV-DakH1279, an isolate from a YFV-infected patient in Senegal in 1965 that has been used previously in non-human primate challenge studies (*12*). Both mAbs had an IC_50_ < 75 ng/mL and IC_90_ < 250 ng/mL against both viruses.

We then performed *in vitro* escape assays to assess mAb-driven viral mutations emerging in the parental YFV-17D infection strain. No envelope protein mutations were detected at any point throughout 10 YFV-17D passages using the two mAbs (data not shown).

### Therapeutic mAb administration protects hamsters from pathogenic YFV infection

Having shown that the two candidate mAbs neutralize pathogenic lab strains and primary isolates of YFV at IC_50_ < 100 ng/mL, we next designed *in vivo* experiments to test the efficacy of mAb administration during acute YFV infection. Syrian golden hamsters were infected with 200 cell culture infectious doses (CCID_50_) of the serially passaged, hamster-adapted YFV-Jimenez strain. Three days post-infection (dpi), we administered a single treatment of either one of our mAb candidates or an SIV-specific, isotype-matched control intraperitoneally at a dose of 20 mg/kg. We monitored disease parameters including survival, viremia, weight change, and serum alanine transaminase (ALT) levels to evaluate the efficacy of the mAb treatments. Both YFV-specific mAbs significantly increased survival, with MBL-YFV-01 administration leading to survival in 10/10 animals (Fig. 1a; p<0.0001). In contrast, 11/15 hamsters treated with the SIV-specific, isotype-matched mAb required humane euthanasia due to severe disease. Next, we assessed YFV infectious titers in the serum of hamsters using an *ex vivo* infectivity assay. We found that only hamsters treated with SIV-specific, isotype-matched mAb had replication-competent YFV in their serum (Fig. 1b). We also measured weights daily and found that hamsters receiving SIV-specific mAb treatment showed significantly reduced weight gain beginning at 6 dpi (Fig. 1c, Extended Data Table 1). Notably, the two control hamsters that survived past 9 dpi also began to gain weight following 12 dpi (Fig. 1c). Finally, we used serum ALT measurements as a biomarker of liver disease. We found significantly elevated serum ALT at 6 dpi only in hamsters with replication-competent YFV in their serum (Fig. 1d). These data show that YFV-specific mAb administration to hamsters following pathogenic YFV challenge can completely reverse disease course and protect hamsters from death.

**Figure 1.**
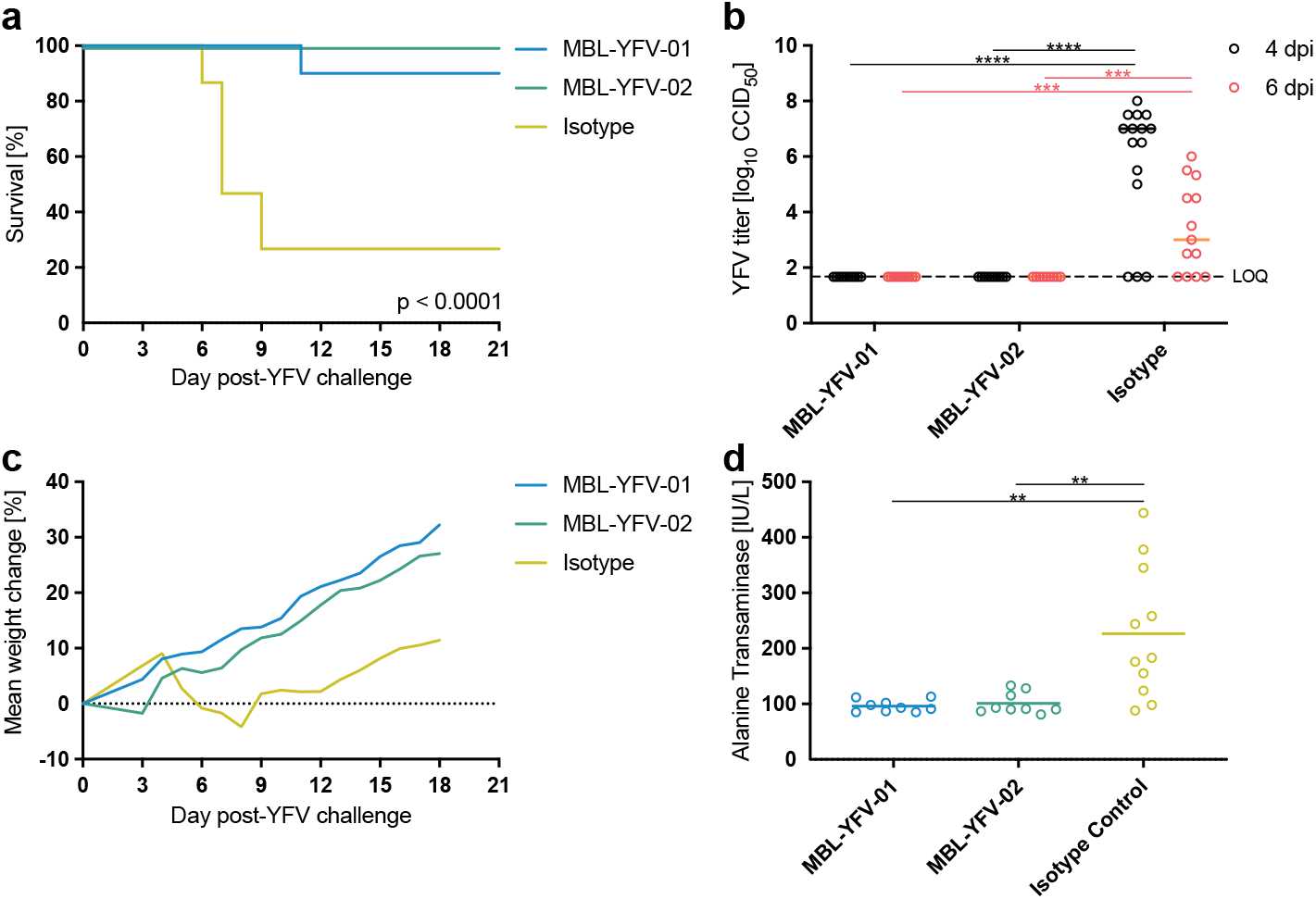
Neutralizing monoclonal antibody administration protects Syrian golden hamsters from lethal YFV infection. a) Kaplan-Meier survival curves of Syrian golden hamsters following challenge with YFV-Jimenez and treatment with YFV-specific or isotype control antibodies. P-value determined by Mantel-Cox test, b) YFV-Jimenez titers in the serum of Syrian golden hamsters at 4 and 6 dpi as measured by an ex vivo infection assay to determine CCID50. P-values determined by one-way ANOVA. c) Longitudinal mean weight change of each hamster treatment group following challenge with YFV-Jimenez. d) Serum alanine transaminase levels of each hamster 6 dpi with YFV-Jimenez. P-values determined by one-way ANOVA.

### Therapeutic mAb administration protects RM from pathogenic YFV infection

Given the success of YFV-specific mAb treatment in hamsters, we next tested MBL-YFV-01 and MBL-YFV-02 in a relevant, preclinical, non-human primate model of pathogenic YFV infection. RM were infected with 1 x 10^3^ CCID_50_ of YFV-DakH1279, a YFV inoculum previously shown to lead to clinical endpoints requiring euthanasia in the majority of RM by 5 dpi (*12*). Two days after YFV challenge, we administered 50 mg/kg of our mAbs intravenously and monitored concentrations of the mAbs in the plasma. Two RMs served as concurrent, untreated controls in the study. Plasma levels of both mAbs reached a mean concentration of 368 μg/mL and were detectable until humane euthanasia at 21 dpi (Fig. 2a). All mAb-treated animals survived for 21 days after infection, while both control animals required humane euthanasia by 5 dpi (Fig. 2b). For statistical comparisons, we included historical animals infected with the same cryopreserved stock and dose of YFV-DakH1279. Five of six required euthanasia (*12*). We found a significant increase in survival of mAb-treated RM compared to controls (p=0.0035). Notably, animals treated with mAb had no detectable YFV in the serum, with the exception of RM 8 at 2 dpi (Fig. 2c). In contrast, both concurrent control RMs had viral loads above 1 x 10^10^ copies/mL at the time of humane euthanasia. Two-way repeated measures ANOVA including the historical controls showed significant reductions in viral loads beginning at 3 dpi (Extended Data Table 1). Next, serum ALT measurements were used as a biomarker of liver disease. We found mildly elevated serum ALT in all mAb-treated animals, with a reduction to baseline levels after mAb treatment. In animals treated with MBL-YFV-01 and MBL-YFV-02 we saw a mean 2-fold and 4-fold ALT increase at 3 dpi, respectively, when compared to baseline (Fig. 2d). A 22-fold increase in peak ALT level was seen in RM 10, while the other concurrent control animal only saw a modest 3-fold increase before meeting other criteria for humane euthanasia (Fig. 2d). YFV RNA was either undetectable or below the level of detection in the livers of mAb-treated animals, but reached high levels in the control RMs (Fig. 2e). Supporting this finding, treated animals had limited gross liver pathology (Fig. 2f), and a single focal point of lymphocytic infiltration in the liver of RM 6 was found by H&E staining (Fig. 2g). In contrast, both concurrent control animals had significant pathological findings at the time of humane euthanasia including severe, diffuse hepatic necrosis and steatosis. Additional pathological findings are included in Extended Data Table 2. Finally, liver staining for YFV RNA from both concurrent controls and a representative animal from each mAb treatment group revealed nearly complete suppression of YFV RNA expression in treated animals, although YFV RNA was detected in the liver of RM 6 (Fig. 2h).

**Figure 2.**
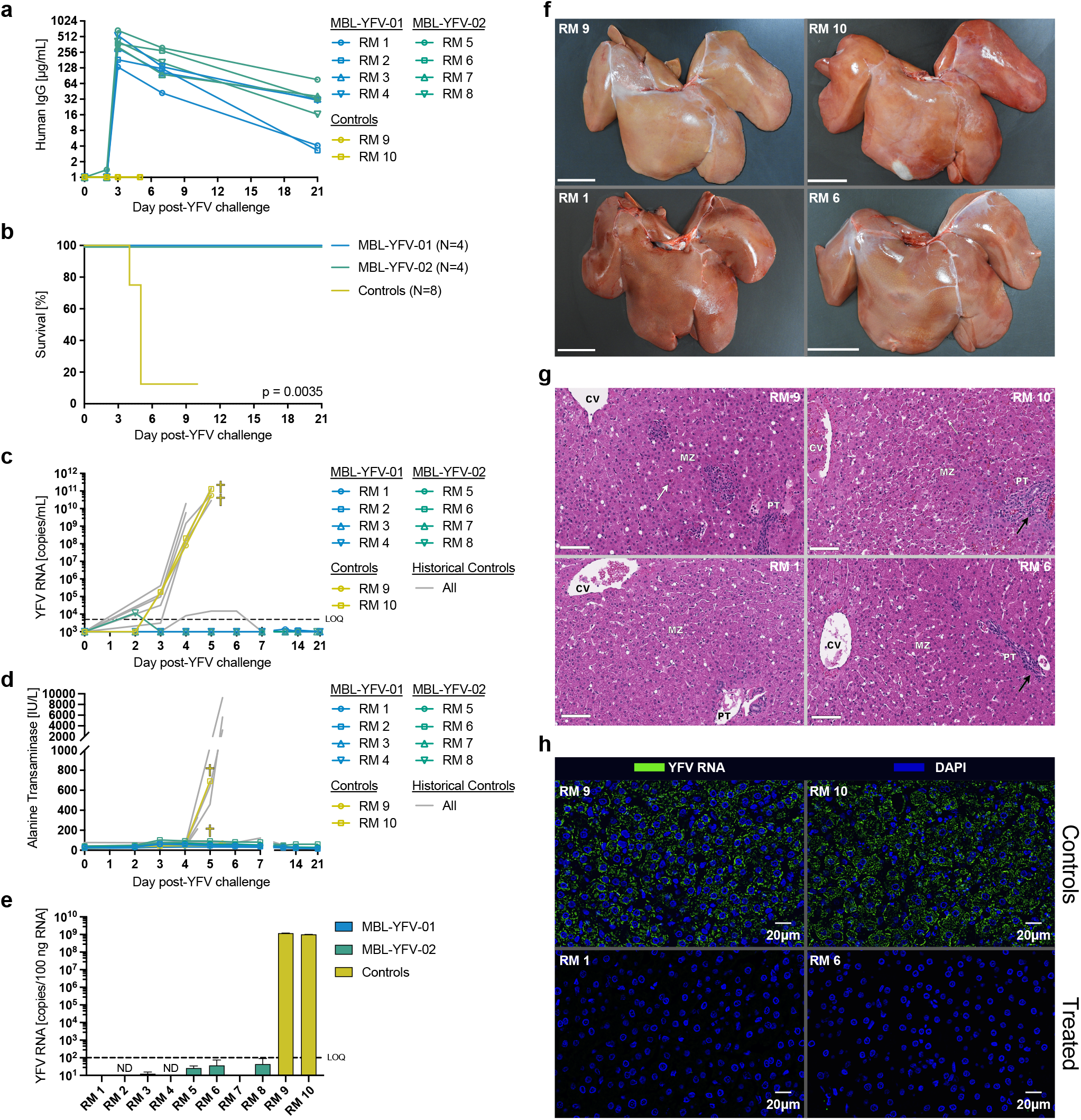
Neutralizing monoclonal antibody administration protects RMs from lethal YFV infection. a) Longitudinal concentration of MBL-YFV-01 and MBL-YFV-02 in the plasma of YFV-DakH1279 challenged RMs. b) Kaplan-Meier survival curves of RMs following challenge with YFV-DakH1279 and treatment with YFV-specific antibodies. Includes historical controls. P-value determined by Mantel-Cox test, c) Longitudinal serum YFV-DakH1279 loads in RMs, including historical controls, d) Longitudinal serum alanine transaminase levels in RMs. e) Quantification of YFV-DakH1279 RNA in the livers of RMs at necropsy, f) Gross pathology of livers of YFV-DakH1279 infected RMs. g) H&E staining of livers of YFV-DakH1279 infected RMs. CV = central vein, PT = portal triad, MZ = medial zone. Black arrows indicate regions of lymphocytic infiltration, h) RNAScope staining of YFV-DakHI279 RNA in the livers of YFV-DakH1279 infected RMs.

## DISCUSSION

There is an urgent need to develop a treatment strategy for YFV infection. In intensive care units, general medical support is the clinical standard (*13*). mAbs are increasingly being used to treat a variety of infectious diseases, and there is precedent that mAbs are effective against viruses that generate very high plasma viral concentrations and cause mortality rates similar to YFV, including Ebola (*14–16*). Here we report on two different mAbs that prevent death in a highly pathogenic non-human primate YFV challenge model.

Our studies demonstrate that a single mAb delivered two days post YFV infection at 50 mg/kg was sufficient to prevent severe disease and death in non-human primates. The ability to utilize a single mAb, rather than a cocktail of mAbs, significantly expedites the pathway to first-in-patient trials. Furthermore, single mAb therapies have recently been shown to be effective therapeutic agents against SARS-CoV-2 infections and have been widely accepted and tolerated in humans (*17, 18*). The RM model system closely resembles human disease progression after YFV infection, particularly in regards to liver dysfunction, hepatocyte necrosis, and the formation of hallmark councilman bodies (*12*). Indeed, YFV infection can lead to liver failure with transplant often being the only available intervention. In the first two months of 2018, surgeons at the University of São Paulo performed five liver transplants in an attempt to save the lives of YFV-infected individuals (*19, 20*). The projected annualized costs in USD for one liver transplant range from $1,427,805 - $2,093,789 which would require only about ten cases to cover the initial costs of manufacturing one of these mAbs (*21*).

During the 2016 outbreak in Brazil, attempts to vaccinate individuals in the affected areas was often too late in preventing infection. Several individuals who became infected and hospitalized with YFV had just received the vaccine (*1*). Indeed, previous studies suggest that the vaccine requires weeks to induce protective neutralizing antibodies, particularly in the elderly (*1, 22*). Additionally, Ab responses generated by the YFV-17D vaccine may not completely neutralize YFV. Therefore, ring-fencing an epidemic by vaccination has limitations in controlling YFV outbreaks, and a readily available therapeutic could help those who fall ill and reduce transmission within these outbreak areas.

The financial burden of Flavivirus infection is estimated to be greater than $2 billion USD per year (*23*) and despite the limited number of recent YFV cases in the USA, the country still remains vulnerable to Flavivirus-induced disease, as shown by the emergence of Dengue and Zika virus (*24–26*). A large portion of the world is also unvaccinated against YFV, putting swaths of people at significant risk. MBL-YFV-01 and MBL-YFV-02 are effective against a variety of YFV strains, and these life-saving medicines could be developed immediately as the first safe and therapeutically efficacious drug for YFV infection.

## MATERIALS AND METHODS

### Antibody expression and purification

Antibody HC and LC variable regions were cloned into human IgG, human IgK, and human IgL backbone constructs. These were transiently transfected and expressed using the Expi293 Expression System (Thermo Fisher). After 6 days, culture supernatants were harvested, spun down, and sterile filtered. mAbs were purified from supernatant using Protein A columns (Cytiva) and eluted all under sterile conditions. Purified mAb used for *in vivo* experiments was then confirmed to be endotoxin-free (Pierce).

### Focus reduction neutralization tests

Focus reduction neutralization tests (FRNTs) were conducted as previously described (*27*). Briefly, mAbs or animal plasma were serially diluted in OptiMEM supplemented with 10% fetal bovine serum and antibiotics. Virus was diluted to a final concentration of approximately 500-1,000 PFU/mL in the same diluent and was added to equal volumes of the diluted plasma and mixed. The virus/plasma mixture was incubated at 37°C for 1 hour. Cell culture medium was removed from 100% confluent monolayer cultures of Vero cells on 96-well plates and 100 μl of the virus/plasma mixture was transferred onto duplicate cell monolayers. Cell monolayers were incubated for 60 min at 37°C with the antibody and virus mixture and then overlaid with 1% methylcellulose in OptiMEM supplemented with 2% FBS, 2 mM glutamine, and 50 μg/mL gentamicin. Samples were incubated at 37 °C for two days after which plaques were visualized by immunoperoxidase staining, and a 50% foci-reduction neutralization titer (FRNT50) was calculated.

### In vitro escape assay

Vero cells were infected with YFV-17D in the presence or absence of mAb. YFV-17D was used at a final MOI of approximately 0.3. Individual purified mAbs were added to the virus at three different concentrations (2, 1, and 0.5 μg/mL) to maximize the opportunity for the virus to select for escape mutants. Virus with antibody or virus and media mixtures were incubated for 1 hr at 37°C before cell adsorption. Cells were infected (1 hr) and then washed and incubated in media with or without mAb supplementation. On the fourth day post infection, 50 μL of the cell-free supernatant was used to infect a new batch of cells. This process was repeated 10 times. From each passage, the remainder of the supernatant was frozen in aliquots for vRNA sequencing. To sequence the envelope from serum virus we isolated viral RNA (Qiagen), synthesized cDNA, and sent for Sanger sequencing using the following primer sets:

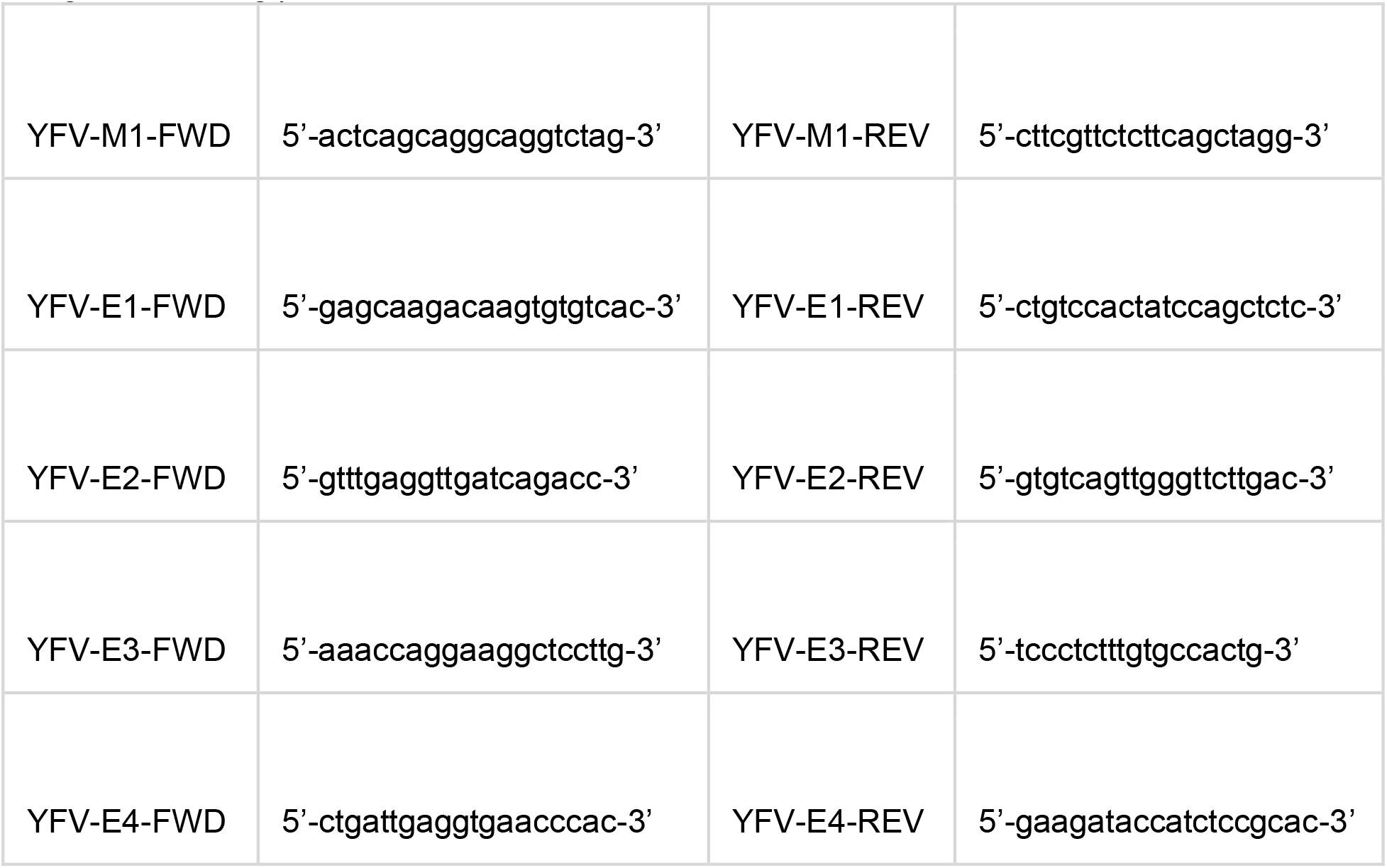

### YFV challenges (hamster)

65 female Syrian golden hamsters (LVG/Lak strain) supplied by Charles River were used. Hamsters were block-randomized by weight to experimental groups and individually marked with ear tags. A challenge dose of 200 CCID_50_ of YFV (Jimenez hamster-adapted strain) per hamster was administered via bilateral i.p. injections in a total volume of 200 μL.

### Infectious cell culture assay (hamster)

Virus titer was quantified using an infectious cell culture assay where a specific volume of either tissue homogenate or serum was added to the first tube of a series of dilution tubes. Serial dilutions were made and added to Vero cells. Ten days later cytopathic effect (CPE) was used to identify the end-point of infection. Four replicates were used to calculate the 50% cell culture infectious doses (CCID_50_) per mL of plasma or gram of tissues.

### Serum aminotransferase assay (hamster)

Serum was collected via ocular sinus bleed 6 dpi. ALT (SGPT) reagent (Teco Diagnostics, Anaheim, CA) was used, and the protocol was altered for use in 96-well plates. Briefly, 50 μL aminotransferase substrate was placed in each well of a 96-well plate, and 15 μL of sample was added at timed intervals. The samples were incubated at 37°C, after which 50 μL color reagent was added to each sample and incubated for 10 minutes as above. A volume of 200 μL of color developer was next added to each well and incubated for 5 minutes. The plate was then read on a spectrophotometer, and aminotransferase concentrations were determined per manufacturer’s instructions.

### YFV challenges and passive antibody administration (RM)

A total of 10 Indian-origin RM were challenged subcutaneously with 1 x 10^3^ CCID_50_ YFV-DakH1279 and assigned to three experimental groups. Intravenous treatment with 50 mg/kg antibody MBL-YFV-01 (N = 4), 50 mg/kg MBL-YFV-02 (N = 4), or no antibody controls (N = 2). Antibodies were infused intravenously at 2 dpi, while controls did not receive any infusions. Animals were cared for at the Oregon National Primate Research Center (ONPRC) with the approval of the Oregon Health & Science University’s Institutional Animal Care and Use Committee using the standards of the NIH Guide for the Care and Use of Laboratory Animals.

### Quantification of delivered huIgG (RM)

Enzyme-linked immunosorbent assay (ELISA) was used to detect delivered human mAbs in plasma. Half-area 96-well costar assay plates (Corning) were coated overnight at 4**°**C with 2 μg/mL of monkey cross-adsorbed goat anti-human IgG (Southern Biotech #2049-01) diluted in phosphate-buffered saline (PBS). Plates were then washed five times with PBS-T (PBS + 0.05% Tween-20) and subsequently blocked with blocking buffer (5% nonfat dry milk in PBS) for 1 hour at 37°C. Standards were prepared in naïve rhesus macaque plasma at a concentration of 50 μg/mL. Heat-inactivated plasma and standards were then serially diluted in blocking buffer. Plates were washed with PBS-T five times, and diluted standards and plasma samples were added to the designated wells. After an hour incubation at 37°C, the plate was washed five times with PBS-T, and detection was carried out with goat anti-human IgG-horseradish peroxidase (HRP) (Southern Biotech, 2045-05) at a 1:10,000 dilution in blocking buffer for 1 hour at 37°C. Plate was then washed five times with PBS-T and developed for 1 minute at room temperature using 3,3’,5,5’-tetramethylbenzidine (TMB) substrate (Southern Biotech). Reactions were stopped with 2 M HCl. Plates were read on the Synergy HTX Multi-Mode Microplate Reader (BioTek) and data was collected using software Gen5 v3.09 at two absorbance wavelengths, 650 nm and 450 nm. The final OD was determined by subtracting OD_650 nm_ from OD_450 nm_.

### YFV-DakH1279 RNA quantification (RM)

Serum viral RNA was extracted from 200 μl using the QiaAmp MinElute Virus Spin kit (Qiagen, Cat# 57704) according the manufacturer’s instructions. Total intracellular DNA and RNA were extracted from the liver tissues. Briefly, samples were snap-frozen in lysing matrix tubes (MP Bio, Cat# 116913050-CF) soaked in 1 mL Trireagent (Molecular Research Center, Catalog# RN190) and homogenized at 4000 rpm for 30 seconds in a Bead BugTM microtube homogenizer (Millipore Sigma, Catalog# Z763713). First, RNA extraction was performed by addition of 1/10th volume bromochloropropane (Sigma Aldrich, Cat# B9673) to the tri-reagent and then vortexed and incubated at room temperature for 5 minutes and then spun at 12,000 x g for 5 minutes to achieve phase separation. 12 μL glycogen (Thermo Fisher, Cat# 10814010) was added to the tubes and the RNA-containing upper aqueous phase was transferred to a fresh tube and placed on ice. Samples for DNA extraction were processed by addition of DNA Backextraction buffer (9.09 g Tris base (Fisher Scientific, Cat# 77-86-1) was added to a 3.75 mL 1M sodium citrate (Fisher Scientific, Cat# BP327-500) solution and 50 mL 6M guanidine thiocyanate (Thermo Fisher, Cat# AM9422), then diluted to a final volume of 75 mL with distilled water) to the sample tubes containing the remaining interphase/organic phase mix after RNA extraction. Samples were vortexed and spun at 12000 x g for 5 minutes to achieve phase separation followed by addition of 12 μL glycogen (Thermo Fisher, Cat# 10814010). The DNA-containing upper aqueous phase was transferred to a fresh tube and placed on ice. Next, both the DNA and RNA samples were treated with isopropanol (Thermo Fisher, Catalog# 383920025) mixed gently by inversion and spun at 15000 x g for 5 minutes at room temperature. The isopropanol was carefully aspirated and 75% ethanol was added to the pellet. The tube was again spun at 15,000 x g for 5 minutes and the ethanol wash repeated a second time. All residual ethanol was removed with a micropipette. The pellets were dried at 37°C on a heat block followed by resuspending the pellets in 100 μL water (RNA) or TE buffer (DNA) (Thermo Fisher, Cat# 12090015). Tubes were placed back in the 37°C heat block and were shaken at 500 rpm for 15 minutes. Tubes were gently vortexed and placed on ice for use in assays. Quantity and integrity of the extracted DNA and RNA were assessed on a NanoDrop 2000 Spectrophotometer (Nanodrop Technologies, Catalog# ND2000).

YFV RNA from both serum and liver was quantified using the TaqPath 1-Step RT-qPCR Master Mix (ThermoFisher Cat# A15299) using primers/probe: YFV_qPCR-F (5’-CACGGGTGTGACAGACTGAAGA-3’), YFV_qPCR-R (5’-CCAGGCCGAACCTGTCAT-3’), and YFV_qPCR-Probe (5’-6FAM-ATGGCGGTG/ZEN/AGTGGAGACGATTG-TAMRA-3’) using an annealing temperature of 60°C. All thermocycling parameters followed exactly to suggested manufacturer’s instructions. All thermocycling and quantification analyses were conducted on an QuantStudio 3 (Applied Biosystems, Catalog# A28567). Quantification was assessed relative to an absolute standard curve using synthesized RNA corresponding to the qPCR target region.

### Serum aminotransferase assay (RM)

Blood was collected into non-anticoagulant, clot activator tubes and spun down at 1860 x g for 10 minutes to separate the serum from the clotted blood. 100 μL of serum was assessed by the Abaxis VETSCAN VS2 Chemistry Analyzer for the quantitative analysis of ALT.

### Pathology assessment (RM)

At the time of necropsy, blood, cerebrospinal fluid, liver, kidney, spleen, hepatic and axillary lymph node(s) with or without a full complement of other tissues were collected. Tissues were fixed in 4% paraformaldehyde for 24 hours and then placed in 70% ethanol for 4-6 days before paraffin embedding. Sections were cut at 5 μm and stained with hematoxylin and eosin on a Leica ST5020 Autostainer and whole slide scanning was performed by a Leica AT2 slide scanner at 20x magnification. Slides were reviewed by two veterinary pathologists using HALO Link software (Indica Labs) and Leica DM 3000 LED microscopes.

### YFV RNA in situ hybridization (RM)

RNA detection was performed using the RNAscope^®^; 2.0 HD Multiplex detection protocol (ACD Life Sciences) followed by Tyramide Signal Amplification (TSA™, Invitrogen) according to manufacturer’s protocol. Liver samples were fixed in 4% paraformaldehyde, processed for paraffin embedding, and cut at 7 μm thickness. Slides were heated at 60°C for 1 hour, dewaxed in xylenes for 10 minutes, and then placed in ethanol 100% for 5 minutes before air drying. Heat-induced epitope retrieval was performed by boiling sections in 1X RNAscope® Pretreat 2 buffer (a citrate buffer [10 nmol/L, pH 6]; ACD Life Science, Catalog# 322000) for 30 minutes, immediately washed in double distilled water, and then dehydrated in 100% ethanol for 5 minutes before air drying. Hydrophobic barrier pen was applied to encircle the section, then the slides were incubated with RNAscope® Protease III (ACD Life Sciences, Catalog# 322337) diluted 1:10 in PBS for 30 minutes at 40°C using a HybEZ hybridization oven (ACD Life Sciences). Slides were rinsed twice with water and then incubated with 1.5% peroxide (Fisher, Catalog# H325-500) for 5 minutes at room temperature. Sections were again rinsed with water and then incubated with pre-warmed customized yellow fever virus (YFV) RNAscope® Probe (V-YFV-ENV-E, ACD Life Sciences, Catalog# 574391) in hybridization buffer A (6X SSC [1XSSC is 0.15 mol/L NaCl, 0.015 mol/L Na-citrate], 25% formamide, 0.2% lithium dodecyl sulfate, blocking reagents) and incubated for 2 hours at 40°C. Slides were washed in wash buffer (0.1X or 0.05X SSC, 0.03% lithium dodecyl sulfate) and incubated with amplification reagents as described in the RNAscope 2.0 HD detection protocol (ACD Life Sciences, Catalog# 322310). Amplifier 1 (2 nmol/L) in hybridization buffer B (20% formamide, 5X SSC, 0.3% lithium dodecyl sulfate, 10% dextran sulfate, blocking reagents) at 40°C for 30 minutes; Amplifier 2 (a proprietary enhancer to boost detection efficiency) at 40°C for 15 minutes; Amplifier 3 (2 nmol/L) in hybridization buffer B at 40°C for 30 minutes; Amplifier 4 (2 nmol/L) in hybridization buffer C (2X SSC, blocking reagents) at 40°C for 15 minutes; Amplifier 5 (a proprietary signal amplifier) at room temperature for 30 minutes; Amplifier 6 (a proprietary secondary signal amplifier) at room temperature for 15 minutes. After each hybridization step, slides were washed with wash buffer three times at room temperature. Before detection, the slides were rinsed one time in 1X TBS Tween-20 (0.05% v/v). Amplification 6 contained alkaline phosphatase (or horseradish peroxidase) labels, which was detected with an Alexa FluorTM 488-conjuagted-tyramide signal amplification (TSA™, Invitrogen, Catalog# B40953) solution for 10 minutes at room temperature. Slides were washed in TBS-T for 10 minutes and then stained with DAPI (5mg/ml, Invitrogen, Catalog# D1306) diluted to 1:10000 in TBS-T for 5 minutes at room temperature. Slides were washed, mounted with SlowFadeTM Gold anti-fade reagent (Invitrogen, Catalog# S36937) and cover-slipped. Images were acquired by scanning the entire tissue at Å~40 magnification using an Olympus VS120 Slide Scanner and pseudocolored using the CellSens™ Dimension Desktop v1.18 software (Olympus).

### Repeated measures statistical analyses

Two-way repeated measures analysis of variance (ANOVA) was used for longitudinal outcome measures to compare yellow fever viral load, ALT, and mean weight change (%) between the antibody-treated and control groups over the study period. In a typical experiment using repeated measures, two measurements taken several time points apart, optimal covariance structure chosen by Bayesian Information Criteria (BIC) was used to account for within subject correlation. Due to right-skewed data, the viral load analysis was performed on log-transformed data. Multiplicity adjusted p-values following Tukey procedures were presented. Statistical significance was determined at the significant alpha level of 0.05. Analyses were performed using SAS version 9.4, specifically PROC MIXED.

## Conflict of Interest

LMW is an author of the patent that describes the monoclonal antibodies in this paper. MJR, DRB, EGK, JBS, and DIW are equity holders and/or employees of Mabloc, LLC.

## Acknowledgments

This work was supported by NIH grants R42 AI155275, P51 OD011092, and by Mabloc, LLC. We would also like to thank the veterinary and animal care staff for their contributions to this study.

