## Extended Data Table 2. for "Therapeutic neutralizing monoclonal antibody administration protects against lethal Yellow Fever infection"

| Animal ID | Pathology |
| --- | --- |
| RM 9 | Hepatic necrosis and steatosis, diffuse, acute, moderate to severe |
| | Icterus, diffuse, minor, renal pelvic fat |
| | Dental calculus, multifocal, chronic, mild |
| | Tension lipidosis, focal, chronic, mild, right medial liver lobe |
| | Adhesions, multifocal, chronic, minimal, caudal lung lobes and diaphragm |
| RM 10 | Hepatic necrosis and steatosis, diffuse, acute to subacute, severe |
| | Dehydration, moderate |
| | Splenomegaly, moderate |
| | Icterus, multifocal, acute, mild, peripancreatic and renal pelvis fat |
| | Lymphadenomegaly, multifocal, mild, peripancreatic lymph nodes |
| | Dental calculus, multifocal, chronic, mild |

**Extended Data Table 2. Gross NHP pathology reports for concurrent control RMs.**
