## Extended Data Table 1. for "Therapeutic neutralizing monoclonal antibody administration protects against lethal Yellow Fever infection"

| Days post-infection | Hamster weight (Fig. 1c) | RM viral loads (Fig. 2c) |
|---|---|---|
| 0 | 1.0000 | 1.0000 |
| 2 | - | 0.9000 |
| 3 | 0.9992 | <0.0001 |
| 4 | 1.0000 | <0.0001 |
| 5 | 0.6578 | <0.0001 |
| 6 | 0.0015 | - |
| 7 | 0.0002 | - |
| 10 | <0.0001 | - |
| 14 | <0.0001 | - |
| 21 | - | - |

**Extended Data Table 1. Two-way repeated measures ANOVA analyses**

**(p-values).**
